# Functional Evaluation of *TRPC6* Missense Variants in Cancer Patients via Molecular Docking Analysis Compared with Patch Clamp Electrophysiology

**DOI:** 10.64898/2026.01.01.697223

**Authors:** Ying Wu, Xiaojing Sun, Joseph S. Reddy, Pooja P. Advani, Nicholas J. Boddicker, James R. Cerhan, Hector R. Villarraga, Samuel J. Asirvatham, Ru-Xing Wang, Hon-Chi Lee, Nadine Norton, Tong Lu

## Abstract

Gain-of-function mutations in the transient receptor potential 6 (TRPC6) channel have recently been recognized as risk factors for both doxorubicin (DOX)-induced cardiomyopathy. Functional evaluation of *TRPC6* missense variants is therefore important for cancer patients undergoing anthracycline treatment. However, traditional electrophysiological methods are labor-intensive and time-consuming. In this study, we compared the functional responses of *TRPC6* missense variants to 1-oleoyl-2-acetyl-sn-glycerol (OAG), a TRPC6 agonist, using molecular docking and patch clamp recording techniques. For the wild-type (WT) TRPC6 structure (PDB ID: 6UZ8), OAG exhibited a binding energy of -4.49 kcal/mol and a dissociation constant (Kd) of 0.511 mM. Twenty *TRPC6* missense variants were identified from cancer patients in the Mayo Clinic database. Of these, fifteen variants had resolvable structures, nine of which displayed increased Kd values and six decreased Kd values compared to WT in molecular docking analysis. Patch clamp recordings revealed that TRPC6 WT and mutant channels were inactive at baseline but were activated upon 50 μM OAG stimulation, except two loss-of-function variants. Moreover, a 24-h treatment with 0.5 μM DOX significantly enhanced OAG-induced channel activation. All three variants identified in patients with heart failure demonstrated gain-of-function properties in both electrophysiological measurements and in-silico predictions. Importantly, the results obtained from molecular docking and patch clamp recordings were strongly correlated, showing an 82% concordance, higher than the predictions from AlphaMissense. These findings indicate that our computational analysis provides a rapid and reliable method for predicting the functional impact of *TRPC6* missense variants, which may aid clinical decision-making in cancer patients receiving chemotherapy.

## INTRODUCTION

Transient receptor potential 6 (TRPC6) is a non-selective cation channel and is ubiquitously expressed in cardiovascular tissues. TRPC6 channels are typically inactive under resting conditions but become activated upon stimulation by diacylglycerol (DAG), a product of phospholipase C (1), and therefore play a critical role in cardiovascular function by regulating intracellular Ca²⁺ signaling (2). Mutations in the *TRPC6* gene significantly affect both focal segmental glomerulosclerosis (FSGS) and cardiovascular disease, underscoring their critical role in Ca²⁺ signaling and cellular homeostasis. In patients with FSGS, both gain-of-function (GOF) and loss-of-function (LOF) mutations in *TRPC6* disrupt podocyte function (3). Beyond the kidney, dysregulated TRPC6 activity in the heart has been implicated in cardiovascular pathologies such as hypertension, atherosclerosis, cardiac hypertrophy, and heart failure (HF) (4–7).

HF is a common cardiovascular disease and is a leading cause of death and hospitalization in the United States (8). Doxorubicin (DOX), one of the most widely used anthracyclines in cancer chemotherapy, is associated with chemotherapy-induced HF. Genome-wide association studies (GWAS) in breast cancer patients have identified intronic and missense variants in *TRPC6* that increase the risk of cardiomyopathy and HF in patients with and without chemotherapy (7, 9–11). Functional characterization of *TRPC6* missense variants is therefore essential to elucidate the molecular mechanisms underlying HF susceptibility, identify therapeutic targets, improve risk stratification, and guide the development of personalized cardioprotective strategies in oncology care. However, traditional electrophysiological approaches are often time-consuming and labor-intensive. In this study, we evaluated *TRPC6* missense variants using 1-oleoyl-2-acetyl-sn-glycerol (OAG), a cell-permeable DAG analog that binds directly to the channel pore region, through both traditional patch-clamp recordings and molecular docking. Our results reveal a strong correlation between OAG responses predicted in silico and those measured by patch clamp. These findings highlight agonist-dependent in-silico analysis as a rapid and effective approach for predicting *TRPC6* variant function, providing a valuable strategy to identify genetic risk variants associated with chemotherapy-induced cardiotoxicity.

## RESULTS

### *TRPC6* variants or mutations identified in cancer patients

Twenty *TRPC6* missense variants were identified in individual patients and mapped across multiple structural domains of the TRPC6 protein, as illustrated in Figure 1A. These included four variants (*P47A*, *P55S*, *R58W*, and *R73H*) located within the unresolved and proximal N-terminal region; six (*R91C*, *S96F*, *N125S*, *D210E*, *I223V*, and *K241N*) in the ankyrin repeat domains; four (*R273K*, *C325S*, *R365H*, and *N338S*) in the linker helices; two (*R399Q* and *A404V*) in the pre-S1 elbow; one (*A681D*) in the pore helix; one (*D798H*) in the unresolved region between the TRP re-entrant and horizontal helices; and two (*Q904R* and *G917A*) in the vertical helix. Ten variants were present in patients who underwent anthracycline-containing chemotherapy. Among them, *N338S* was previously reported in a single patient who developed heart failure post-treatment and *A404V*, a polymorphism with a frequency of ∼12% in the general population, was significantly associated with decline in left ventricular ejection fraction (LVEF) in breast cancer patients treated with doxorubicin and with HF not related to anthracycline (7). Notably, *G917A* variant was found in a lymphoma patient who had not received chemotherapy but still developed HF and arrhythmia. Cardiac phenotypes of patients carrying *TRPC6* missense variants are summarized in Table 1.

**Figure 1.**
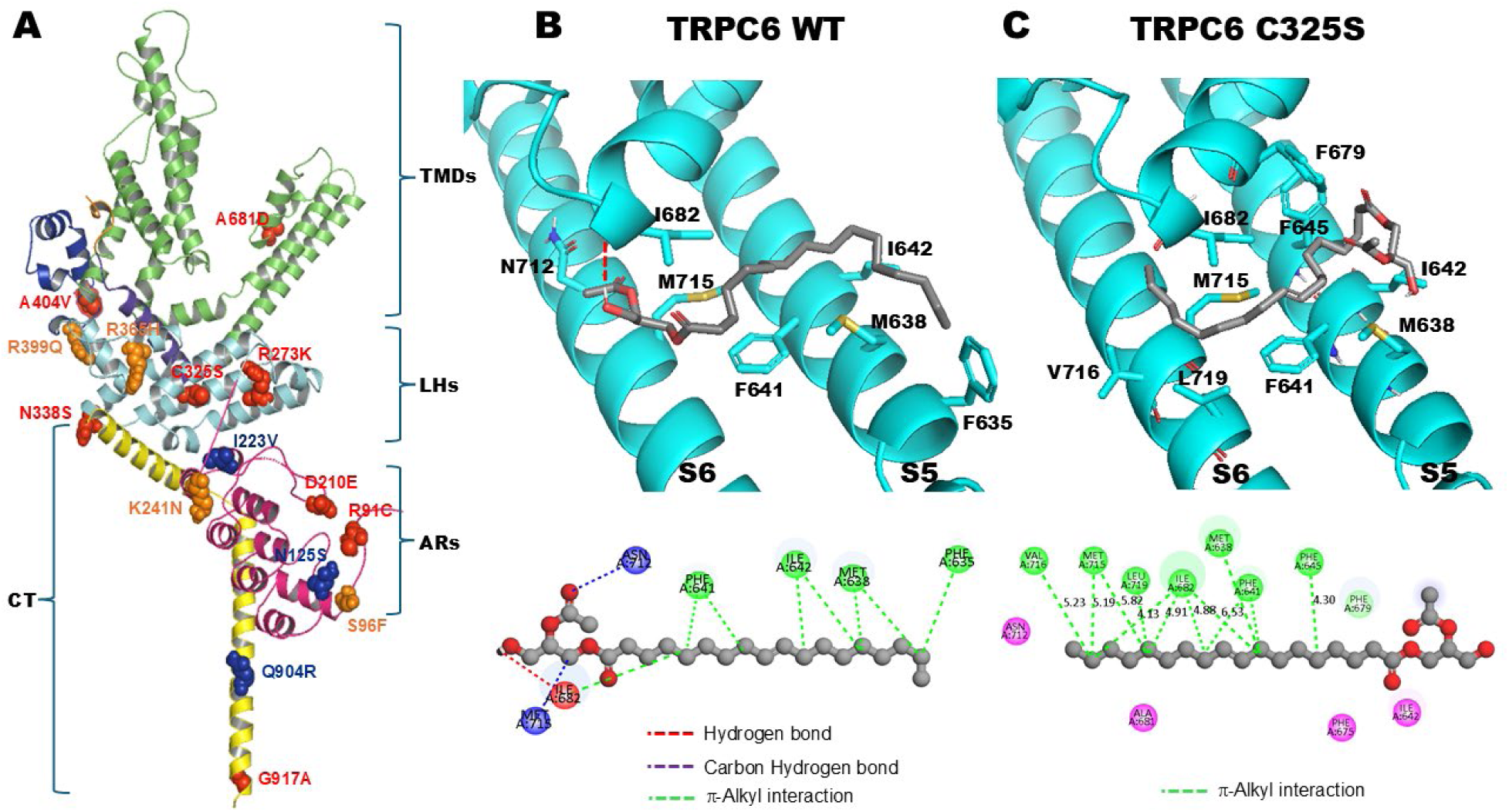
Evaluation of *TRPC6* missense variant function by molecular docking analysis **A:** The cryo-EM structure of human TRPC6 (PDB ID: 6UZ8) and the locations of missense variants tested in this study are shown. Each TRPC6 subunit contains cytoplasmic N- and C-terminus (CT) and six transmembrane domains (TMDs, green), which include a voltage-sensing domain (S1-S4) and a pore-forming domain (S5-S6) with a pore helix. The N-terminus contains four ankyrin repeats (ARs, magenta), followed by nine linker helices (LHs, grey). The C-terminus comprises a vertical helix (VH, yellow) and a horizontal helix (HH, yellow). Several key structural elements are characteristic of TRPC6, including the pre-S1 elbow (violet), a three-helix region preceding S1, and the highly conserved transient receptor potential helix (TRPH, light blue), followed by the TRP re-entry helix that connects S6 to the HH (brown). Based on in-silico analysis, gain-of-function *TRPC6* variants (*R91C*, *D210E*, *R273K*, *C325S*, *N338S*, *A404V*, *A681D*, and *G917A*) are highlighted in red, silent variants (*S96F*, *K241N*, *R365H*, and *R339Q*) in orange, and loss-of-function variants (*N125S*, *I223V*, and *Q904R*) in blue. **B:** Molecular interaction between TRPC6 and OAG. In TRPC6 WT, OAG binds to a groove formed by the S5 and S6 domains, as well as the pore-helix, with the binding energy of -4.5 kcal/mol and apparent Kd value of 0.51 mmol/L. Specifically, I682 residue in the pore-helix forms a hydrogen bond with the oleic acid molecule and interacts hydrophobically with the glycerol backbone of OAG. Two additional carbon-hydrogen bonds are formed between N712 and M715 residues in the S6 region of TRPC6 WT and the oleic acid molecule of OAG. The glycerol backbone of OAG forms multiple alkyl/π-alkyl interactions with F635, M638, F641, and I642 residues in S5 of TRPC6 WT. **C:** Molecular interaction between TRPC6 C325S and OAG. TRPC6 C325S exhibits interactions with OAG similar to WT channel but involving more residues. Notably, F625 and F679 in the S5 segment, along with V716 in the S6, contribute to these interactions by providing additional hydrophobic and other non-covalent contacts. These conformational changes lead to a decreased binding energy (-4.92 kcal/mol) and a reduced apparent Kd (0.249 mM) for the OAG-TRPC6 C325S interaction.

**Table 1:** Cardiac Manifestations of *TRPC6* Missense Variants in Patients with Cancer

| **Variant** | **Frequency in Genome AD** | **Dataset** | **Cancer Type** | **Anthracycline (y/n)** | **Cardiotoxicity (y/n)** | **Variant Function** |
| --- | --- | --- | --- | --- | --- | --- |
| P47A | 0.00006209 | MER | lymphoma | y | n | unknown |
| P55S | 0.000004405 | Mayo Clinic Cardiotoxicity registry | breast cancer | y | n | GOF |
| R58W | 0.001229 | MER | lymphoma | n | n | unknown |
| R73H | 0.00003242 | MER | lymphoma | y | n | unknown |
| R91C | 0.00005081 | MER | lymphoma | n | n | unknown |
| S96F | not present | MER | lymphoma | y | n | silent |
| N125S | 0.0003414 | MER | lymphoma | n | n | LOF |
| D210E | 0.000002478 | MER | lymphoma | y | n | unknown |
| I223V | 0.0001035 | MER | lymphoma | y | n | LOF |
| K241N | 0.000006195 | MER | lymphoma | n | n | silent |
| R273K | 6.195e-7 | MER | lymphoma | y | n | GOF |
| C325S | 6.196e-7 | MER | lymphoma | y | n | GOF |
| N338S | 0.00001239 | N9831 clinical trial | breast cancer | y | y<br>heart failure (7) | GOF |
| R365H | 0.00004152 | MER | lymphoma | n | n | silent |
| R399Q | 0.00005390 | MER | lymphoma | n | n | LOF/silent* |
| A404V | 0.1115 | present in all datasets | breast cancer,<br>lymphoma | y | y<br>heart failure (7) | GOF |
| A681D | not present | MER | breast cancer | y | n | GOF |
| D798H | 0.0004895 | MER | lymphoma | n | n | unknown |
| Q904R | 0.00006196 | MER | lymphoma | n | n | silent/LOF* |
| G917A | 0.000004957 | MER | lymphoma | n | y | GOF |

AD: aggregation database. MER: molecular epidemiology resource. GOF: gain-of-function. LOF: loss-of-function. \*: the differences between the molecular docking prediction and electrophysiological assessment (in-silico analysis/patch-clamp recordings).

### Molecular docking analysis of TRPC6 mutant structures with OAG

All mutants, except those related to unresolved structures, were generated using the cryo-electron microscopy (cryo-EM) structure of human TRPC6 (PDB ID: 6UZ8), followed by energy minimization prior to molecular docking analysis. As we have previously reported, OAG binds to a pocket formed by the pore-helix, the transmembrane domain S5 and S6 within the same subunit of TRPC6 protein (12, 13). Specifically, residues F635, M638, F641, I642, I682, N712, and N715 in TRPC6 wild type (WT) interact with OAG through hydrogen bonds and multiple hydrophobic interactions (Figure 1B). The OAG binding energy was calculated at -4.49 kcal/mol with a computer-derived apparent dissociation constant (Kd) of 0.511 mM. Despite variations in residues, all TRPC6 mutant structures retain a similar OAG-binding site. For instance, the interaction between OAG and the TRPC6 C325S involves additional residues (M638, F641, I642, F645, F679, I682, N712, M715, and L719), resulting in enhanced non-covalent hydrophobic interactions. This leads to a lower OAG binding energy (-4.92 kcal/mol) and a higher Kd value (0.247 mM), indicating a favorable binding affinity between OAG and TRPC6 C325S (Figure 1C).

Given that the apparent Kd values for TRPC6 channels typically varied within 0.1 mM across three independent simulations, we categorized the TRPC6 mutant channels into three groups based on their Kd values: <0.4 mM, between 0.4 and 0.6 mM, and >0.6 mM. Among these variants, eight (*R91C, D210E, R273K, C325S, N338S, A404V, A681D*, and *G917A*) exhibited apparent Kd values below 0.4 mM; four variants (*S96F, K241N, R365H*, and *R399Q*) had Kd values between 0.4 and 0.6 mM; and three (*N125S, I223V*, and *Q904R*) showed Kd values greater than 0.6 mM. Table 2 presents detailed information on the binding energy, computer-derived Kd values, number of hydrogen bonds, and the amino acid residues involved in the interactions between TRPC6 mutant channels and OAG, as well as the pathogenic scores of AlphaMissense.

**Table 2:**
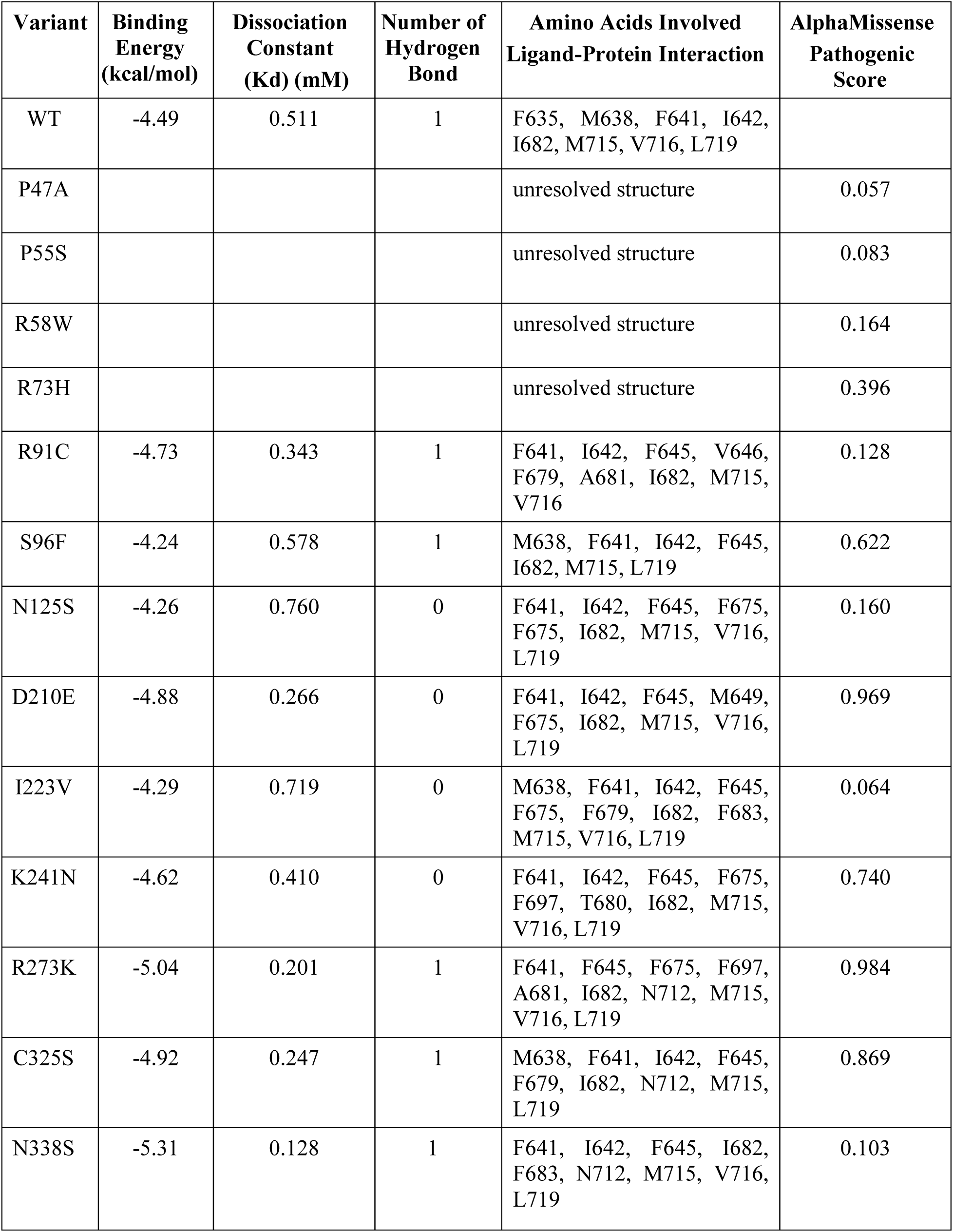
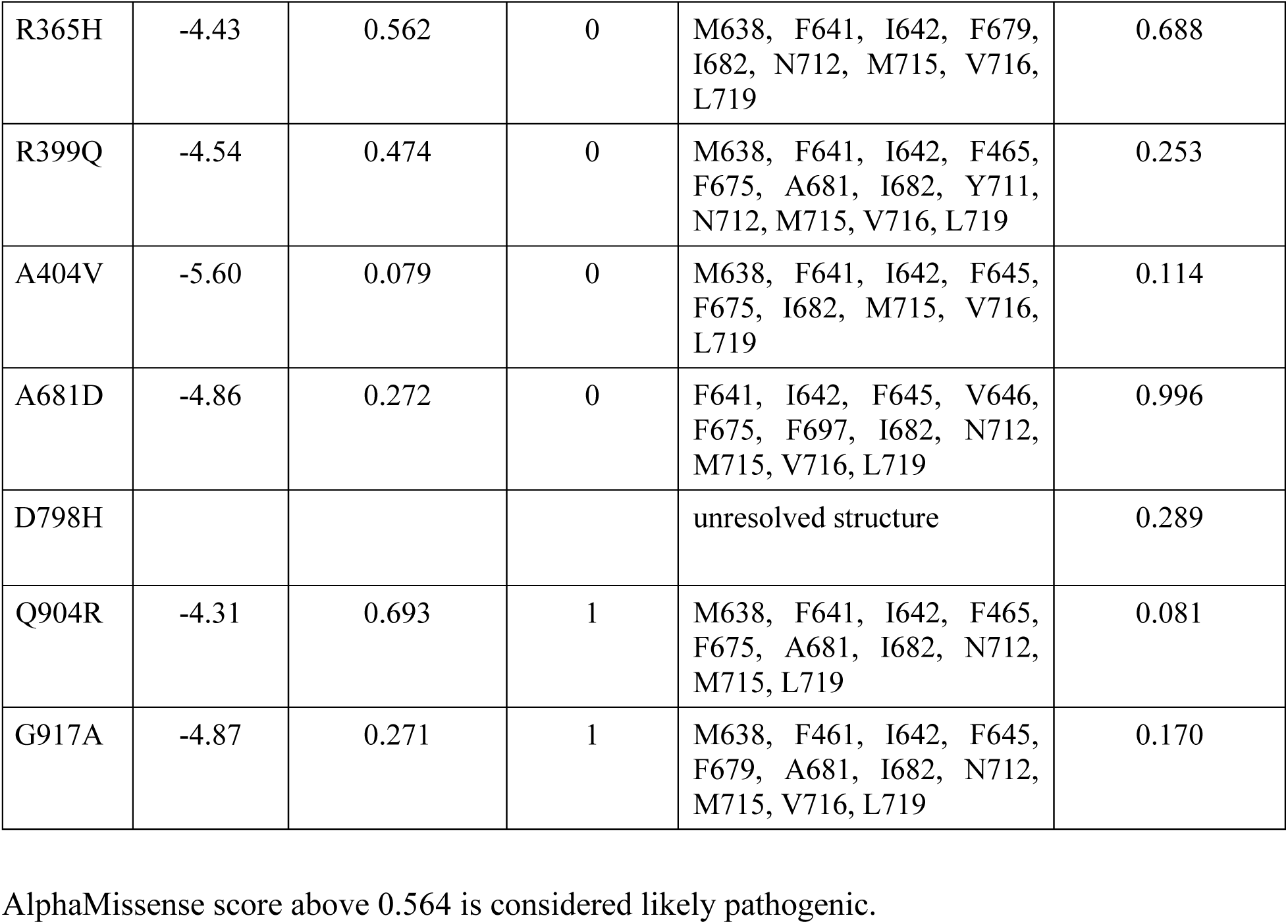
Functional Prediction by Molecular Docking and AlphaMissense Scoring

### Electrophysiological examination of TRPC6 mutant channel activities

To further validate our computational modeling results, we performed patch-clamp experiments to assess OAG-mediated activation of TRPC6 mutant channels, including three mutants with Kd values between 0.4 and 0.6 mM (S96F, R365H, R399Q), four with Kd values below 0.4 mM (R273K, C325S, A404V, G917A), and three with Kd values above 0.6 mM (N125S, I223V, Q904R). Additionally, we examined one mutant with an unresolved structure (P55S), all in comparison to WT controls.

Figure 2 presents representative current traces from TRPC6 WT channels (Kd: 0.511 mM) and the S96F mutant (Kd: 0.578 mM) before and after application of 50 µM OAG. In WT channels, OAG elicited a 1.8-fold increase in inward current densities, from -6.35±1.38 to -11.33±2.19 pA/pF at -100 mV (n=10, p<0.05), and a 4.1-fold increase in outward current densities, from 10.76±2.51 to 44.43±8.97 pA/pF at +100 mV (n=10, p<0.05). In contrast, the S96F mutant showed a significantly reduced response. Inward current densities increased 2.4-fold from -3.54±0.60 to - 8.60±1.42 pA/pF at -100 mV (n=11, p<0.05 vs. WT), and outward current densities increased 3.8-fold from 13.27 ± 1.65 to 50.20 ± 5.10 pA/pF at +100 mV (n=11, p<0.05 vs. WT).

**Figure 2.**
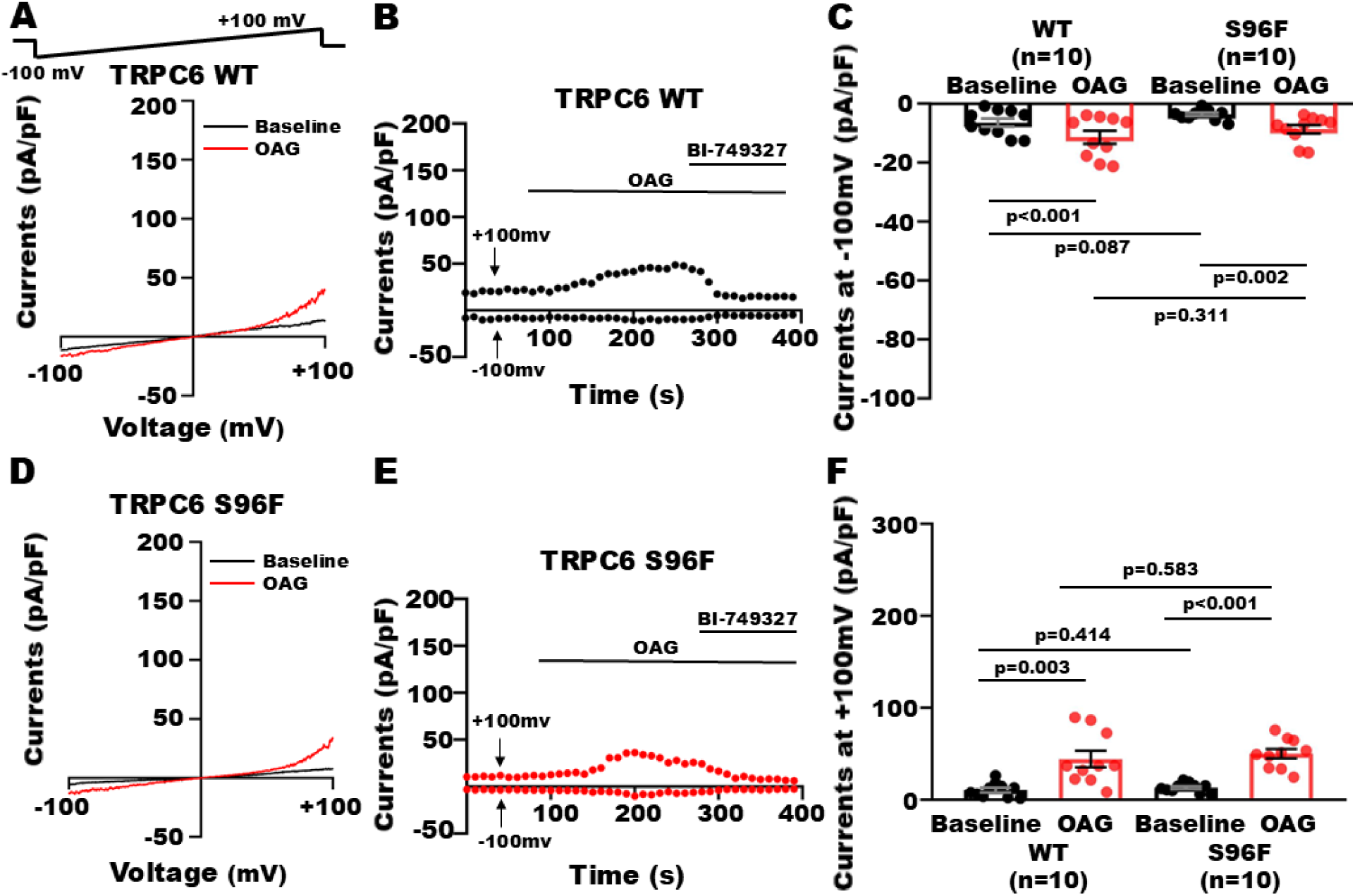
Effect of OAG on TRPC6 S96F channels at baseline. **A and D:** Whole-cell currents were recorded from HEK293 cells expressing TRPC6 WT and S96F mutant using a voltage-ramp protocol from -100 mV to +100 mV over 100 ms at a holding potential of -60 mV. Application of OAG (50 μM) enhanced inward and outward currents of TRPC6 S96F channels as much as that of WT controls. **B and E:** The time course of OAG effect on inward (measured at -100 mV) and outward current densities (measured at +100 mV) of TRPC6 WT and S96F channels. The effects of OAG on TRPC6 channels were abolished by superfusion with BI-749327 (1 μM). **C and F**: Group results with individual data points and statistical significance are shown in the bar graphs (Paired t-test and Student’s t-test).

Following a 24-h treatment with 0.5 μM DOX, OAG-induced responses in WT channels were significantly enhanced. Inward current densities increased from -2.87±0.51 to to -24.68±4.78 pA/pF at -100 mV (n=8, p<0.05 vs. baseline and OAG without DOX), while outward current densities augmented from 10.14±0.92 to 83.75±11.46 pA/pF at +100 mV (n=8, p<0.05 vs. baseline and OAG without DOX). DOX treatment also modestly potentiated OAG responses in S96F channels. Inward current densities increased from -3.10±0.36 to -10.10±1.03 pA/pF at -100 mV (n=11, p<0.05 vs. baseline and WT+DOX), and outward current densities enhanced from 15.91±2.50 to 72.97±5.97 pA/pF at +100 mV (n=11, p<0.05 vs. baseline and WT+DOX), representing 3.3-fold and 4.6-fold increases, respectively (Figure 3). These results indicate that the S96F is a silent mutation.

**Figure 3.**
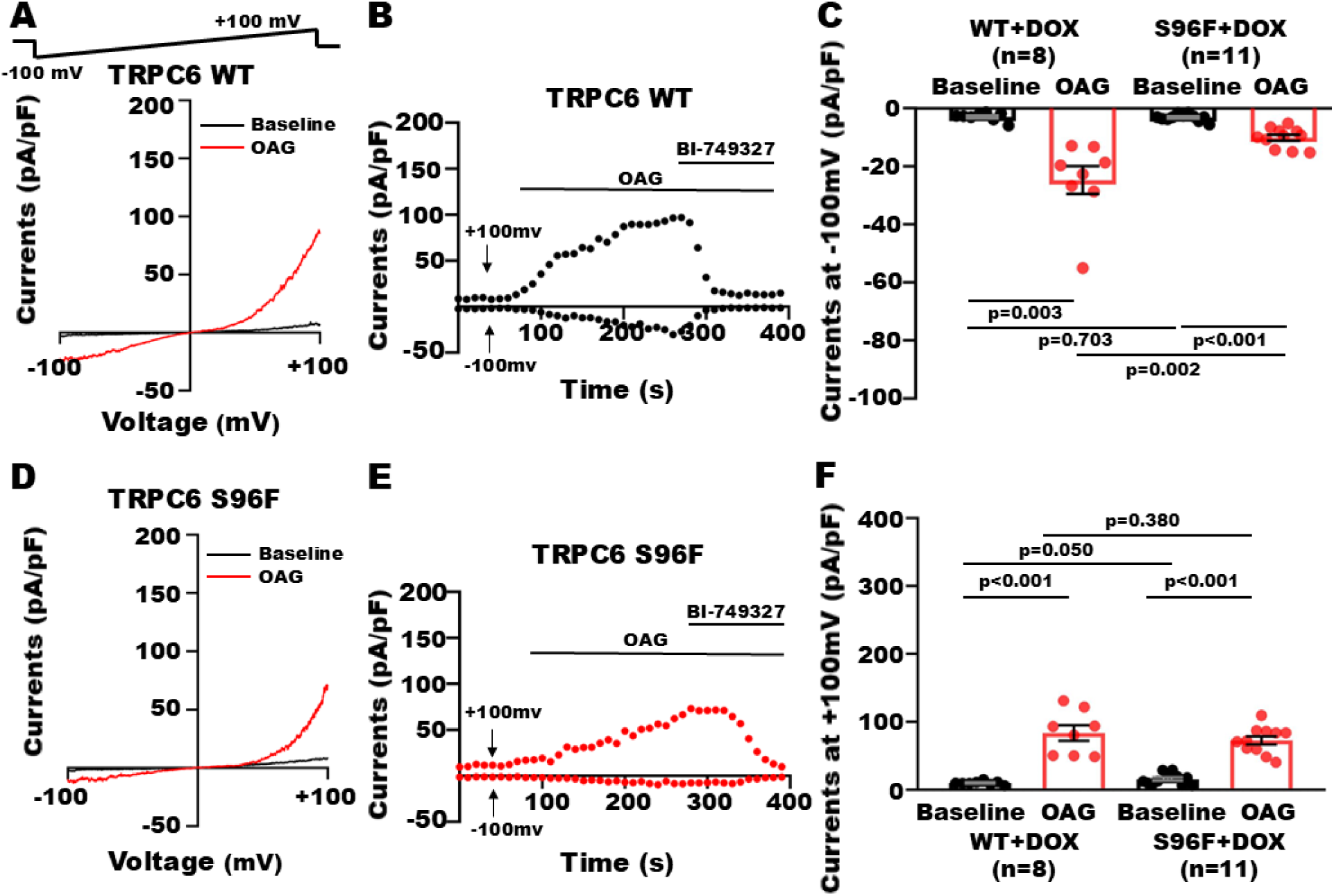
Effect of OAG on TRPC6 S96F channels after DOX treatment. **A and D:** Whole-cell currents were elicited from HEK293 cells 24 h after transfection with TRPC6 WT and TRPC6 S96F cDNAs, followed by an additional incubation with 0.5 μM DOX for 24 h, using a voltage-ramp protocol from -100 mV to +100 mV over 100 ms at a holding potential of - 60 mV. Incubation with DOX enhanced the effects of 50 μmol/L OAG on both inward and outward current densities of TRPC6 WT and S96F channels, with no significant difference between them. **B: and E:** The time course of OAG effect on inward (measured at -100 mV) and outward current densities (measured at +100 mV) of TRPC6 WT and S96F channels after incubation with DOX. The effects of OAF were inhibited by exposure to BI-749327. **C and F:** Group results with individual data points and statistical significance are shown in the bar graphs (Paired t-test, Student’s t-test or Mann-Whitney test).

Figure 4 displays representative current traces from WT and C325S mutant channels (Kd: 0.247 mM) in response to 50 μM OAG. In WT channels, OAG increased inward current densities from -5.31±1.32 to -16.5±2.83 pA/pF at -100 mV (n=11, p<0.001), and outward current densities from 15.02±2.83 to 61.35±10.31 pA/pF at +100 mV (n=11, p<0.001), corresponding to 3.1-fold and 4.1-fold increases, respectively. In contrast, the C325S mutant displayed a markedly enhanced response: inward current densities increased 8.3-fold from -3.38 ± 0.60 to -28.17±3.59 pA/pF at -100 mV (n=12, p<0.001 vs. WT), and outward current densities augmented 7.6-fold from 13.54±1.95 to 103.32±8.93 pA/pF at +100 mV (n=12, p<0.001 vs. WT). The baseline current densities did not differ significantly between WT and C325S channels.

**Figure 4.**
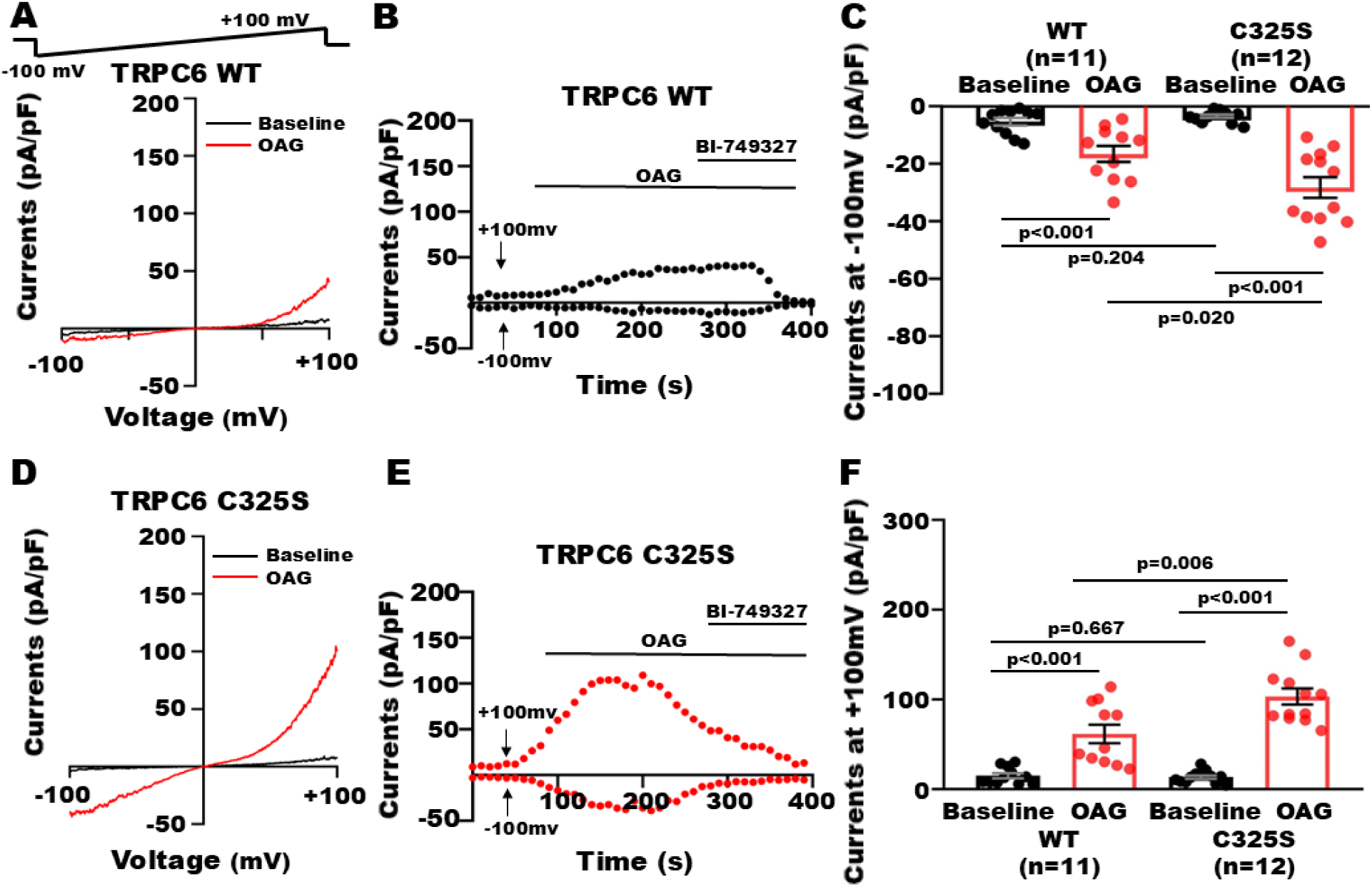
Effect of OAG on TRPC6 C325S channels at baseline. **A and D:** Whole-cell currents were elicited by a voltage-ramp protocol from -100 mV to +100 mV over 100 ms from a holding potential of -60 mV in HEK293 cells 48 h after transfection with TRPC6 WT and TRPC6 C325S cDNAs. At baseline, the C325S mutant channel has similar inward and outward current densities, compared to WT. Application of OAG (50 μM) robustly increased the inward- and outward-currents of TRPC6 WT, and the OAG effects were significantly greater in TRPC6 S325S channels. **B and E**: The time course of OAG effect on inward (measured at -100 mV) and outward current densities (measured at +100 mV) of TRPC6 WT and S325C channels. The effects of OAG on TRPC6 channels were abolished by application of BI-749327 (1 μM), a highly selective TRPC6 inhibitor. **C and F:** Group results with individual data points and statistical significance are shown in the bar graphs (Paired t-test and Student’s t-test).

Treatment with 0.5 μM DOX for 24 h further enhanced OAG-induced currents in both WT and C325S channels. In WT channels, inward current densities increased from -5.41±1.14 to - 25.46±7.57 pA/pF at -100 mV (n=8, p<0.05), and outward current densities elevated from 7.70±1.16 to 75.44±19.21 pA/pF at +100 mV (n=8, p<0.05 vs. baseline and OAG without DOX). In C325S channels, DOX treatment led to a dramatic increase in OAG responses: inward current densities rose 19.9-fold from -1.63±0.22 to -32.46±5.93 pA/pF at -100 mV (n=11, p<0.001 vs. baseline and WT+DOX), and outward current densities increased 15.8-fold from 10.14±1.48 to 159.94±21.26 pA/pF at +100 mV (n=11, p<0.001 vs. baseline and WT+DOX) (Figure 5). Hence, the C325S is a GOF mutant.

**Figure 5.**
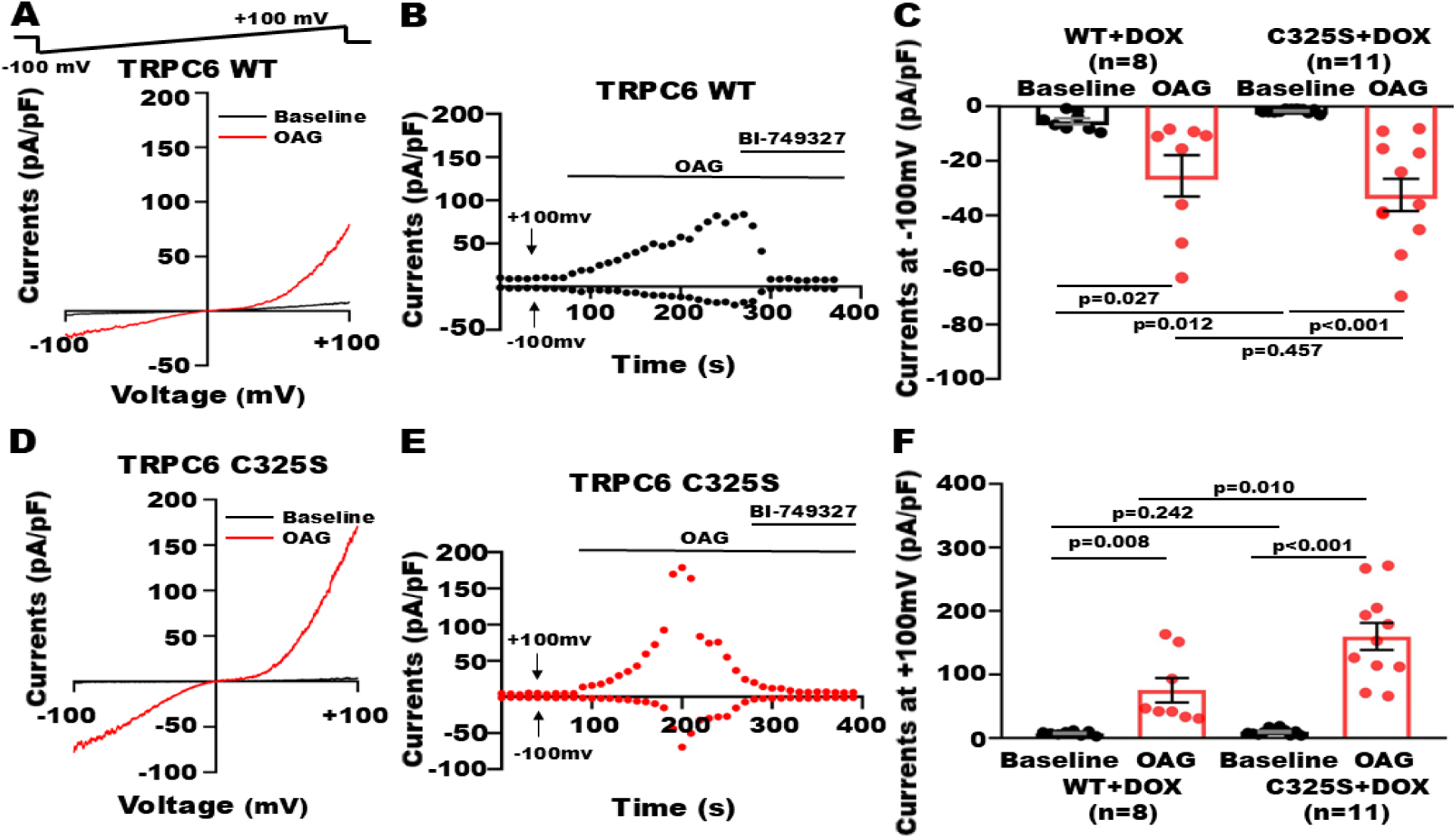
Effects of OAG-mediated TRPC6 WT and C325S channel activation after DOX treatment. **A and D:** Whole-cell currents were elicited by a voltage-ramp protocol from -100 mV to +100 mV over 100 ms from a holding potential of -60 mV in HEK293 cells 24 h after transfection with TRPC6 WT and TRPC6 C325S cDNAs, followed by an additional incubation with DOX (0.5 μmol/L) for 24 h. Treatment with DOX further potentiated the OAG (50 μM) effects on outward current increase but not on inward current increase in TRPC6 C325S channels, compared to the effects of OAG on TRPC6 WT channels. **B and E:** The time course of OAG effect on inward (measured at -100 mV) and outward current densities (measured at +100 mV) of TRPC6 WT and C325S channels after incubation with DOX. The effects of OAG on TRPC6 channels were abolished by superfusion with BI-749327 (1 μM). **C and F**: Group results with individual data points and statistical significance are shown in the bar graphs (Paired t-test, Student’s t-test or Mann-Whitney test).

Similar experimental strategies were applied to N125S mutant channels (Kd=0.76 mM). In WT controls, OAG significantly enhanced current densities: inward current densities increased from -2.91±1.10 to -11.38±1.84 pA/pF at -100 mV (n=11, p<0.05), and outward current densities rose from 7.96±1.79 to 37.40±4.82 pA/pF at +100 mV (n=11, p<0.001), corresponding to 3.9-fold and 4.7-fold increases, respectively. In contrast, OAG-induced responses were markedly attenuated in N125S mutant channels. Inward current densities showed only a modest 1.1-fold increase, from -0.98±0.10 to -1.08±0.05 pA/pF at -100 mV (n=11, p=0.412 vs. WT), while outward current densities increased 2.0-fold, from 5.09±0.60 to 10.10±2.73 pA/pF at +100 mV (n=11, p=0.079 vs. WT) (Figure 6).

**Figure 6.**
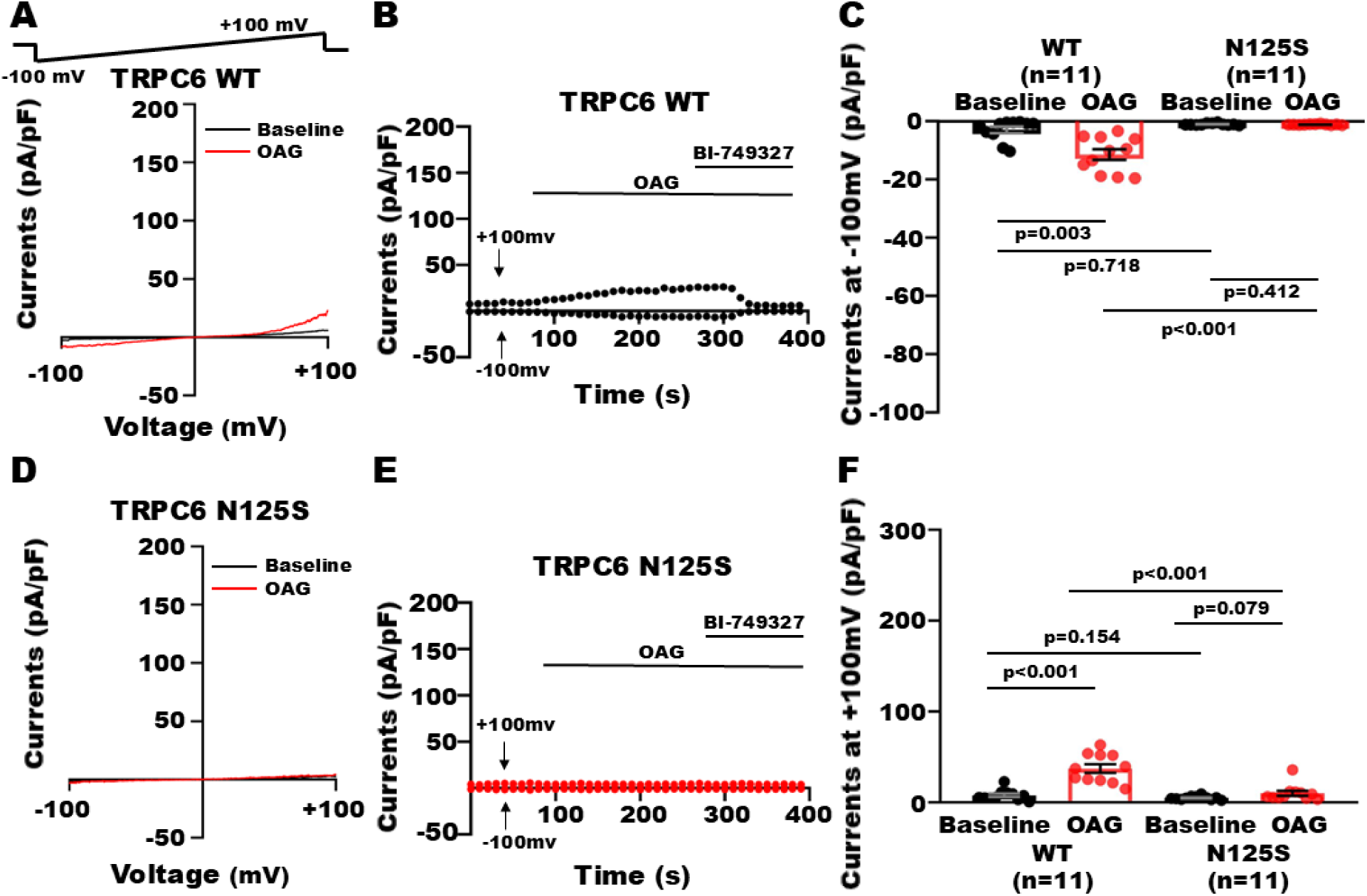
Effect of OAG on TRPC6 N125S channels at baseline. **A and D:** Whole-cell currents were recorded in HEK293 cells 48 h after transfection with TRPC6 WT or TRPC6 N125S cDNAs, using a voltage-ramp protocol from -100 mV to +100 mV over 100 ms, with a holding potential of -60 mV. At baseline, the N125S mutant channel is inactive and has no response to OAG (50 μM), compared to WT control. **B and E**: The time course of OAG effect on inward (measured at -100 mV) and outward current densities (measured at +100 mV) of TRPC6 WT and N125S channels. **C and F:** Group results with individual data points and statistical significance are shown in the bar graphs (Paired t-test, Student’s t-test or Mann-Whitney test).

Following a 24-h treatment with DOX, WT channels exhibited significantly amplified responses to OAG. Inward current densities increased from -2.15±1.01 to -21.28±3.77 pA/pF at - 100 mV (n=7, p<0.05 vs. baseline and OAG without DOX), while outward current densities enhanced from 8.53±2.19 to 72.30±14.27 pA/pF at +100 mV (n=7, p<0.05 vs. baseline and OAG without DOX). However, on N125S channels, DOX treatment had a very small but significant effect on channel activation after application of OAG. There was no change in inward current densities from -1.38±0.25 to -1.28±0.17 pA/pF at -100 mV (n=11, p=0.622 vs. baseline and WT+DOX), but slight increase in outward current densities from 4.71±1.07 to 8.67±2.04 pA/pF at +100 mV (n=11, p<0.05 vs. baseline and WT+DOX) (Figure 7). Our results suggest that the N125S is an LOF mutant.

**Figure 7.**
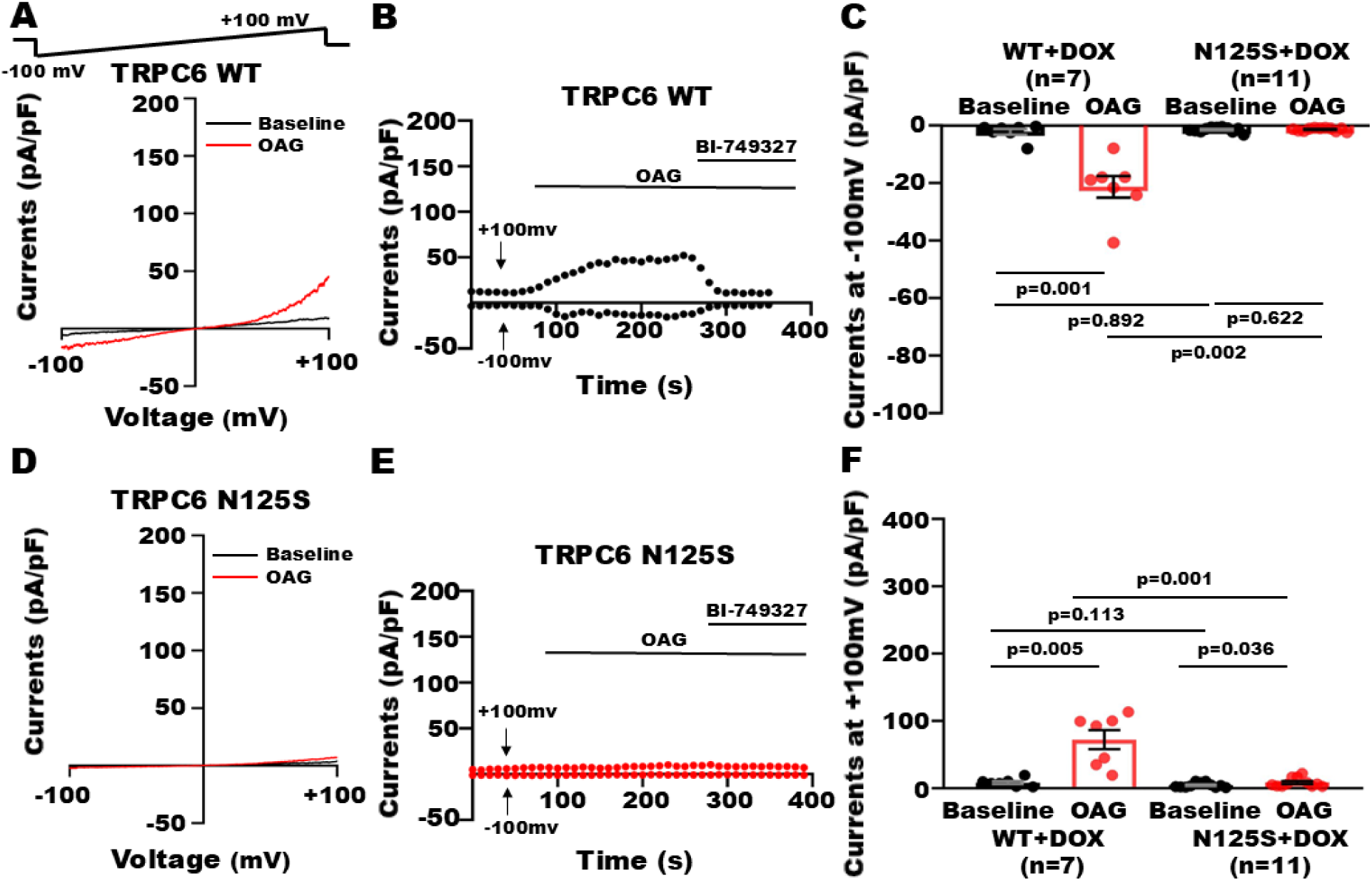
Effect of OAG on TRPC6 N125S channels after DOX treatment. **A and D:** Whole-cell currents were recorded in HEK293 cells 24 h after transfection with TRPC6 WT and TRPC6 N125S cDNAs, followed by an additional incubation with DOX (0.5 μM) for 24 h, using a voltage-ramp protocol from -100 mV to +100 mV over 100 ms with a holding potential of -60 mV. OAG (50 μM) had no effect on inward current densities, but slightly and significantly increased outward current densities in TRPC6 N125S mutant channels after DOX treatment. **B and E:** The time course of OAG effect on inward (measured at -100 mV) and outward current densities (measured at +100 mV) of TRPC6 WT and N125S channels after incubation with DOX. **C and F**: Group results with individual data points and statistical significance are shown in the bar graphs (Paired t-test, Student’s t-test or Mann-Whitney test).

Additionally, we evaluated the function of *P55S* mutant in an unresolved region of the 6UZ8 structure and confirmed that it is a GOF mutant. Without DOX treatment, OAG increased inward current densities of P55S channels by 3.2-fold, from a baseline of -7.30 ± 1.08 pA/pF to - 23.55 ± 4.17 pA/pF at -100 mV (n=10, p<0.05) and outward current densities by 5.2-fold, from 18.55 ± 4.6 pA/pF to 96.42 ± 20.83 pA/pF at +100 mV (n=10, p<0.05). Following a 24-h incubation with DOX, the effects of OAG were further amplified, showing a 6.8-fold increase in inward current densities from -8.27 ± 0.57 pA/pF to -56.04 ± 10.82 pA/pF at -100 mV (n=10, p<0.05) and a 11.7-fold enhancement in outward current densities from 13.97 ± 3.41 pA/pF to 163.23 ± 24.03 pA/pF at +100 mV (n=10, p<0.001).

### Correlation between patch clamp examination and in-silico analysis

Figure 8 summarizes the effects of OAG on inward and outward current densities of TRPC6 WT and mutant channels in the absence and presence of DOX treatment. To evaluate the relationship between OAG-induced increases in TRPC6 current densities in patch-clamp recording and the computer-derived Kd values from molecular docking, we performed a linear regression analysis. The Kd values (ranked from low to high) were plotted against the fold increases in the inward current densities at -100 mV and the outward current densities at +100 mV induced by OAG in the absence (Figure 8A) and presence of DOX treatment (Figure 8B). The analysis revealed a strong negative correlation between the apparent Kd values and the magnitudes of current increments, with Pearson correlation coefficient (r) ranging from -0.73 to -0.90 (p=0.01 ∼ <0.001). This reciprocal relationship was consistently observed both at baseline and DOX-treatment conditions, reinforcing the predictive value of molecular docking in estimating TRPC6 channel responsiveness to OAG activation.

**Figure 8:**
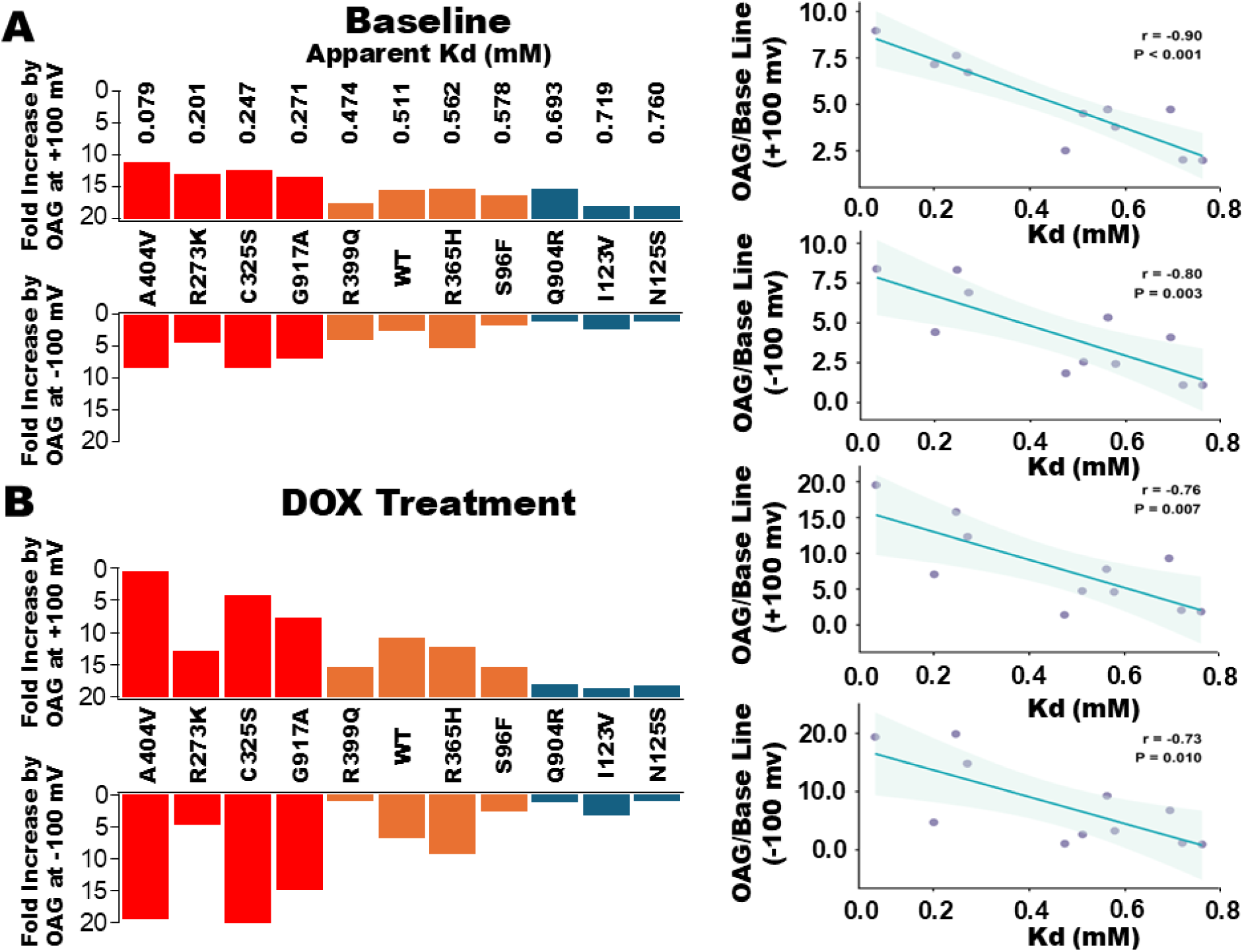
Correlation between molecular docking and patch-clamp recording approaches in characterizing *TRPC6* missense variant function. Based on the computer-derived dissociation constant (Kd), four mutants with Kd values below 0.4 mM, three with Kd values between 0.4 and 0.6 mM, and three with Kd values above 0.6 mM were selected for electrophysiological assessments in comparison to WT control. The bar graphs illustrate OAG-induced fold changes in inward current densities (measured at -100 mV) and outward current densities (measured at +100 mV) of TRPC6 WT and mutant channels under baseline conditions (**A**) and after a 24-h DOX treatment (**B**), arranged in order of increasing Kd values. Strong negative correlations were observed between apparent Kd and fold increases in current densities with Pearson correlation coefficients (r) ranging from -0.73 to -0.90.

## DISSCUSSION

TRPC6 channels are the key mediators of Ca^2+^ influx in response to mechanical and chemical stimuli, playing a critical role in maintaining normal renal and cardiovascular function. In the kidney, both GOF and LOF *TRPC6* mutations have been linked to the development of FSGS (3, 14, 15). GOF *TRPC6* variants have also been associated with HF in patients, regardless of chemotherapy exposure (7, 9, 10). In contrast, LOF *TRPC6* variants have shown protective effects in animal models, conferring resistance to pathological cardiac hypertrophy and preserving cardiac function (16). These findings underscore the pivotal role of TRPC6 in renal and cardiovascular pathophysiology and highlight its potential as a therapeutic target.

Traditional methods for examining TRPC6 mutant channel activity are often time-consuming and labor-intensive, requiring site-directed mutagenesis, patch-clamp recording, and intracellular Ca²⁺ imaging in heterologous expression systems. In this study, we compared the functional impact of *TRPC6* missense variants identified in cancer patients using molecular docking analysis and patch-clamp electrophysiology. All TRPC6 channels displayed very low baseline current densities, comparable to WT controls, but responded robustly to OAG activation except for two LOF mutants. These results suggest that TRPC6 channels remain largely inactive under resting conditions and require external stimulation for functional activation, consistent with previous reports (13, 16).

Three out of seven GOF *TRPC6* variants were observed in individuals with cancer-related cardiotoxicity or in non-cancer patients with HF (7, 13). Notably, the rare N338S variant was identified in a breast cancer patient who developed HF following doxorubicin and trastuzumab treatment (7). The common *A404V* variant has been associated with chemotherapy-induced declines in LVEF and HF in early breast cancer patients treated with DOX, as well as in non-cancer patients with HF and FSGS (7, 17, 18). Additionally, the *G917A* variant was identified in a lymphoma patient who did not receive anthracyclines but developed HF and arrhythmia following alternative cancer therapies. The fact that not all individuals with GOF variants developed anthracycline-induced HF may be due to low penetrance, potentially influenced by individual genetic, environmental, and lifestyle factors (19). Interestingly, none of LOF variants were observed in patients with HF or other cardiovascular disorders in the datasets in this study. Based on our results from the current study and a previous report (13), our molecular docking analysis was consistent with patch clamp assessments in 82.8% of cases, except for the *R339Q* and *Q904R* variants. Specifically, *R339Q* displayed an LOF phenotype electrophysiologically but was classified as silent by molecular docking, whereas *Q904R* appeared functionally silent in electrophysiology but was predicted as LOF in silico (Figure 8A). Notably, neither variant was associated with HF. A strong reciprocal correlation was observed between the computer-derived Kd values and fold-increases in TRPC6 channel current densities upon OAG activation, both in the absence and presence of DOX, indicating strong concordance between molecular docking predictions and electrophysiological measurements. These findings demonstrate that computational modeling of TRPC6-OAG interactions provides an efficient approach for evaluating the functional impact of *TRPC6* missense variants.

Recently, artificial intelligence has emerged as a powerful tool for predicting the pathogenicity of missense variants in clinical genetics (20, 21). Tools such as AlphaMissense, combined with clinical databases like ClinVar, have been applied to predict the pathogenicity of variants of unknown significance (22). AlphaMissense has analyzed all 71 million possible missense variants in the human genome, classifying 89% of them as either likely benign (57%) or likely pathogenic (32%) (22). Recently, pathogenicity assessment of long QT syndrome-associated ion channel variants using traditional patch-clamp techniques and AlphaMissense predictions revealed strong concordance, with 80% sensitivity and 76% specificity (23).

We compared *TRPC6* missense variant predictions between our molecular docking and AlphaMissense scoring. Of the seven GOF variants identified, three (*R273K, C325S, and A682D*; 42.9%) exhibited high pathogenic potential with AlphaMissense scores of 0.984, 0.869, and 0.996, respectively, exceeding the pathogenicity threshold of 0.564 (22). However, patients carrying these variants did not develop anthracycline-related HF or other cardiovascular complications (Table 1). Conversely, all LOF variants (*N125S, I223V, R399Q,* and *Q904R*; 100%) were predicted as benign, exhibiting higher Kd values for OAG interaction and lower AlphaMissense scores. Interestingly, two of three functionally silent mutants (*R365H and K241N*; 66.7%) were classified as likely pathogenic, with AlphaMissense scores of 0.740 and 0.688, respectively. Surprisingly, three *TRPC6* variants (*N338S, A404V*, and *G917A*), previously identified in patients with HF and confirmed by electrophysiology and molecular docking to have GOF effects, were classified as benign by AlphaMissense, with low scores of 0.013, 0.114, and 0.170 (Table 2). In addition, *N125S* and *A404V* have been causally associated with FSGS (18). These discrepancies may occur because AlphaMissense focuses solely on predicted structural conformational changes in *TRPC6* missense variants and does not include functional activation mechanisms such as post-translational modulation and agonist-driven stimulation. They may also reflect that TRPC6-induced cardiac malfunction is affected by personal genetic, environmental, and lifestyle factors that current computational models cannot account for (19).

Our study has several limitations. First, because of the high conservation of *TRPC6* coding sequence and the rarity of missense variants, the number of cancer patients carrying *TRPC6* missense variants who received anthracycline therapy was relatively small. Larger cohorts will be necessary to more accurately assess the sensitivity and specificity of our in-silico analysis. Second, the reliability of molecular docking is highly dependent on the quality of available three-dimensional TRPC6 structures. Although high-resolution cryo-EM structures (5YX9, 6UZ8, 6UZA, and 7A6U) have improved our understanding of TRPC6 architecture, some regions remain unresolved. Additionally, AlphaFold-predicted models provide relatively complete structures, however, they predict only a single static conformation and lack integration of ligands, cofactors, or post-translational modifications, which may reduce accuracy in disordered or flexible regions and protein complex predictions (24). These structural limitations may, in turn, affect the precision of computational predictions.

In summary, reliable evaluation of *TRPC6* missense variants critically depends on agonist-induced activation, and molecular docking provides a powerful, rapid approach to predict their functional consequences, potentially guiding precision therapy for cancer patients receiving anthracyclines.

## MATERIALS and METHODS

### Resources of human TRPC6 variants or mutations

Twenty *TRPC6* variants were identified in patients with lymphoma from the Molecular Epidemiology Resource (MER) (IRB #09-001987) (25) and in patients with breast cancer from our in-house cardiotoxicity registry at Mayo Clinic Florida (IRB #19-002566 and IRB #22-001501) and from the completed N9831 clinical trial (IRB #15-2789) (7, 9, 26). Exome sequencing of the MER cohort was performed by Regeneron Genetic Center as previously described (27). Exome sequencing of patients in the cardiotoxicity registry was performed by the Mayo Clinic Medical Genome Facility. DNA samples from the subset of patients in the N9831 clinical trial with chemotherapy-related HF were sequenced by Sanger sequencing for TRPC6 exons only as previously described (7).

### Molecular docking analysis of TRPC6 structures with OAG

Molecular docking between TRPC6 structure and OAG interaction was performed as previously described (13). Briefly, the cryo-EM structure of human TRPC6 (PDB ID: 6UZ8) was obtained from the National Center of Biotechnology Information (NCBI) protein database, and single TRPC6 mutant was generated using the PyMOL v3.1.3.1 software (Schrödinger, LLC., New York, NY). The two-dimensional structure of OAG (PubChem CID: 6504449) was downloaded from PubChem database and then converted into a three-dimensional structure using Avogadro v1.99.0 software (Avogadro Chemistry, University of Pittsburgh, Pittsburgh, PA). Energy minimization of TRPC6 WT and mutant structures were conducted using Swiss-pdb viewer v4.1 software (Swiss Institute of Bioinformatics, Switzerland). Molecular docking was performed using AutoDockTools v1.5.7 (Molecular Graphics Laboratory, The Scripps Research Institute, La Jolla, CA) and analyzed using BIOVIA Discovery Studio Visualizer (Dassault Systèmes SE, Corp., France). The OAG binding energy and computer-derived equilibrium dissociation constant (Kd) values were calculated as the average of three independent repeats.

### Cell culture, TRPC6 mutagenesis and expression

HEK293 cells were purchased from American Type Culture Collection/ATCC (Manassas, VA) and maintained in Dulbecco’s Modified Eagle Medium (DMEM, Sigma-Aldrich, Inc., St. Louis, MO) supplemented with 10% fetal bovine serum and 100 μg/ml penicillin-streptomycin. Human TRPC6 WT cDNA in pcDNA3.1+C-(K) DYK (Clone ID: OHu23592) was obtained from GenScript, Inc. (Piscataway, NJ). TRPC6 mutagenesis was carried out using the QuikChange Site-Directed Mutagenesis Kit (Catalog #200517, Agilent Technologies, Inc., Santa Clara, CA). The orientation and accuracy of TRPC6 mutations was confirmed by DNA sequencing. Recombinant TRPC6 cDNAs and green fluorescent protein (GFP) cDNAs were co-transfected into HEK293 cells using Effectene Transfection Kit (Cat. #301425, Qiagen, Inc., Gaithersburg, MD) as we have previously described (13). Transfected cells were detected by observing GFP expression under ultraviolet microscope (Olympus, IX70, Olympus America, Inc., Savage, MN).

### Whole-cell patch clamp recordings

Forty-eight h after transfection, TRPC6 channel currents were evoked using a 100-ms voltage-ramp protocol from -100 mV to +100 mV at a holding potential of -60 mV as previously described (13). The bath solution contained (in mM): NaCl 140.0, KCl 4.0, CaCl_2_ 2.0, MgCl_2_ 1.0, HEPES 10.0, and glucose 5.0, at pH 7.4. The pipette solution contained (in mM): Cs^+^-aspartate 145.0, MgCl_2_ 2.0, CaCl_2_ 0.3, EGTA 10.0, and HEPES 10.0, at pH 7.35. Total currents were continuously recorded from HEK293 cells at baseline and application of 50 μM OAG. Following the maximum effect of OAG, 1 μM of BI-749327 (a highly selective TRPC6 inhibitor) was perfused. TRPC6 currents, defined as BI-749327-sensitive components, were obtained by subtracting the total currents from residual currents in the presence of BI-749327. Data acquisition was performed using Clampex 10.7 and subsequently analyzed with Clampfit 10.7 software (Molecular Devices, LLC, San Jose, CA).

### AlphaMissense-based pathogenicity prediction of *TRPC6* mutations

The pathogenicity of human *TRPC6* variants (TRPC6_HUMAN, Q9Y210, ENST00000344327.8) was assessed using the AlphaMissense online tool (https://alphamissense.hegelab.org/) (28), with scores above 0.564 considered as having pathogenic potential (22).

### Chemicals

OAG (Cat. # 495414) was obtained from Cayman Chemical Company, Inc. (Ann Arbor, MI). BI-749327 (Cat. # HY-111925) was purchased from MedChemExpress, LLC. (Monmouth Junction, NJ). Unless otherwise mentioned, all chemicals were purchased from Sigma-Aldrich, Inc.

### Statistical analysis

Data are presented as means ± standard error of the mean (SEM). Statistical analyses were performed using R programming language (www.r-project.org), as previously described (29). For normally distributed data, paired t-tests were used to compare means before and after treatment. One-way ANOVA with Holm’s post-hoc test was employed to assess differences among groups. For non-normally distributed data, the Wilcoxon signed-rank test was applied for two-group comparisons, and the Kruskal-Wallis H test with Dunn’s post-hoc test was used for multiple-group comparisons. Pearson’s correlation coefficients (r) between variables were calculated using the “cor” function in R. Statistical significance was defined as p<0.05.

## ACKNOWLEDGEMENTS

We would like to thank the following teams and individuals for their contributions:

## Mayo Clinic Project Generation

**Leadership Team:** James R. Cerhan, Fergus J. Couch, and Janet E. Olson.

Statistical Genetics and Bioinformatics Team: Nicholas B. Larson and Zachary S. Fredericksen

**Laboratory Operations:** Mine Cicek.

**Management Team:** Lisa K. Colborn, Andrew J. Danielsen, Jonathan J. Harrington, and Jennifer

1. M. Kushwaha.

## Mayo Clinic Regeneron Genetics Center Project Generation

**Management & Leadership Team:** Aris Baras, Gonçalo Abecasis, Adolfo Ferrando, Giovanni Coppola, Andrew Deubler, Luca A. Lotta, John D. Overton, Jeffrey G. Reid, Alan Shuldiner, Katherine Siminovitch, Jason Portnoy, Marcus B. Jones, Lyndon Mitnaul, Alison Fenney, Jonathan Marchini, Manuel Allen Revez Ferreira, Maya Ghoussaini, Mona Nafde, William Salerno, Cristen Willer, Lourdes Crane.

**Sequencing & Lab Operations:** John D. Overton, Christina Beechert, Erin Fuller, Laura M. Cremona, Eugene Kalyuskin, Hang Du, Caitlin Forsythe, Zhenhua Gu, Kristy Guevara, Michael Lattari, Alexander Lopez, Kia Manoochehri, Prathyusha Challa, Manasi Pradhan, Raymond Reynoso, Ricardo Schiavo, Maria Sotiropoulos Padilla, Chenggu Wang, Sarah E. Wolf, Hang Du, Kristy Guevara.

**Genome Informatics & Data Engineering:** Jeffrey G. Reid, Mona Nafde, Manan Goyal, George Mitra, Sanjay Sreeram, Rouel Lanche, Vrushali Mahajan, Sai Lakshmi Vasireddy, Gisu Eom, Krishna Pawan Punuru, Sujit Gokhale, Benjamin Sultan, Pooja Mule, Mudasar Sarwar, Muhammad Aqeel, Xiaodong Bai, Lance Zhang, Sean O’Keeffe, Razvan Panea, Evan Edelstein, Ayesha Rasool, William Salerno, Evan K. Maxwell, Boris Boutkov, Alexander Gorovits, Ju Guan, Lukas Habegger, Alicia Hawes, Olga Krasheninina, Samantha Zarate, Adam J. Mansfield, Lukas Habegger.

**Analytical Genetics and Data Science**: Gonçalo Abecasis, Manuel Allen Revez Ferreira, Joshua Backman, Kathy Burch, Adrian Campos, Liron Ganel, Sheila Gaynor, Benjamin Geraghty, Arkopravo Ghosh, Salvador Romero Martinez, Christopher Gillies, Lauren Gurski, Eric Jorgenson, Tyler Joseph, Michael Kessler, Jack Kosmicki, Adam Locke, Priyanka Nakka, Jonathan Marchini, Karl Landheer, Olivier Delaneau, Maya Ghoussaini, Anthony Marcketta, Joelle Mbatchou, Arden Moscati, Anita Pandit, Jonathan Ross, Carlo Sidore, Eli Stahl, Timothy Thornton, Sailaja Vedantam, Rujin Wang, Kuan-Han Wu, Bin Ye, Blair Zhang, Andrey Ziyatdinov, Yuxin Zou, Jingning Zhang, Kyoko Watanabe, Mira Tang, Frank Wendt, Suganthi Balasubramanian, Suying Bao, Kathie Sun, Chuanyi Zhang, Sean Yu, Aaron Zhang, David Corrigan, Dhruv Shidhaye, Chen Wang, Keyrun Adhikari, Alexander Lachmann.

**Research Program Management & Strategic Initiatives:** Marcus B. Jones, Michelle G. LeBlanc, Nadia Rana, Jennifer Rico-Varela, Jaimee Hernandez, Larizbeth Romero, Ashley Paynter.

**Senior Partnerships & Business Operations:** Randi Schwartz, Lourdes Crane, Alison Fenney, Jody Hankins, Anna Han, Samuel Hart, Ryan Smith.

**Business Operations & Administrative Coordinators:** Ann Perez-Beals, Gina Solari, Johannie Rivera-Picart, Michelle Pagan, Sunilbe Siceron.

This work is dedicated to the memory of Professor Wen-Ping Jiang, whose mentorship, insight, and constant encouragement profoundly influenced the beginning of my research journey.

## AUTHOR CONTRIBUTION

Y.W., X.S., J.S.R., P.P.A., N.J.B. and J.R.C. contributed to data collection and analysis. R.W., H.L., and N.N. interpreted the results and revised the manuscript. T.L. designed the study, analyzed results, and wrote the manuscript.

## Abbreviations

Cryo-EM: cryo-electron microscopy
DAG: diacylglycerol
DOX: doxorubicin
FSGS: focal segmental glomerulosclerosis
GFP: green fluorescent protein
GOF: gain-of-function
GWAS: genome-wide association study
HEK: human embryonic kidney
HF: heart failure
Kd: equilibrium dissociation constant
LOF: loss-of-function
LVEF: left ventricular ejection fraction
MER: molecular epidemiology resource
OAG: 1-oleoyl acetyl-sn-glycerol
pA/pF: picoamperes per picofarad
TRPC6: transient receptor potential canonical 6

## DATA AVAILABILITY

All data generated during this study are available within the article. Further any extra information regarding the experimental details and data will be available upon reasonable request.

